# Age-related Facilitation of Bimanual Finger Behaviors by Haptic Augmentation

**DOI:** 10.1101/2024.04.29.591778

**Authors:** Matti Itkonen, Niko Lappalainen, Hana Vrzakova, Roman Bednarik

## Abstract

Proficient use of multiple fingers across both hands enables complex interactions with the world. Developing dexterity for delicate bimanual tasks can take weeks to years to master. This study examines whether a simple mechanical abstraction can enhance bimanual response times to visual stimuli using index fingers and thumbs. A plate placed over the buttons was introduced to shift task conceptualization—rather than pressing buttons, subjects rocked the plate over them. Following this mechanical augmentation, we observed improved bimanual performance in middle-aged participants but no effect on unimanual actions. Conversely, younger adults showed no performance improvement. A control group of middle-aged participants confirmed that the observed improvements resulted from the augmentation. We hypothesize that highly similar afferent sensory signals from the fingers of both hands alter the motor control strategy. This work provides a foundation for further research into the diverse mechanisms of multi-fingered bimanual behavior in both abstract and practical contexts.

## 1 Introduction

Humans rely on the collaborative use of both hands for daily activities (Bailey, Klaesner, & Lang, 2015). Fine manipulation tasks, such as using a knife and fork (Grannen, Wu, Belkhale, & Sadigh, 2022), playing musical instruments (Chang et al., 2014), or performing surgery (Koskinen, Huotarinen, Elomaa, Zheng, & Bednarik, 2022), often require years of practice to achieve bimanual proficiency. Similarly, operators of articulated cranes and other complex man-machine control systems must develop task-specific bimanual coordination to manage multiple degrees of freedom effectively (Suzuki, Takase, Pan, Ishikawa, & Furuta, 2008).

Mastering complex fine manipulation presents two primary challenges: independent control of individual fingers (Latash & Zatsiorsky, 2009) and independent coordination of both hands (Janczyk, Schneider, & Hesse, 2022). Complete control over every finger joint is anatomically impossible and exceeds cognitive motor control limits (Santello, Baud-Bovy, & Jörntell, 2013). Additionally, coordinating two limbs simultaneously leads to competition for neural resources within the central nervous system (CNS) (Sardar, Yeo, Allsop, & Punt, 2023; Janczyk et al., 2022). This phenomenon, known as *bimanual interference* (Ivry, Diedrichsen, Spencer, Hazeltine, & Semjen, 2004), manifests as unintended spatial and temporal coupling of the hands (Haken, Kelso, & Bunz, 1985).

Acquiring bimanual proficiency is a gradual process (Fitts & Posner, 1967), requiring days to weeks for neural adaptations that integrate new behaviors (Pascual-Leone, Amedi, Fregni, & Merabet, 2005). Learned behaviors form internal models that associate actions with predicted sensory outcomes, allowing rapid adaptation to changing internal and external conditions (Wolpert & Kawato, 1998).

Beyond gradual adaptation, sensory feedback modulation (Kovacs & Shea, 2010; Alnajjar et al., 2020) and task conceptualization (Sartori, Spoto, Gatti, & Straulino, 2020; Banakou, Kishore, & Slater, 2018; Franz, Zelaznik, Swinnen, & Walter, 2001) can immediately influence motor behavior. This study investigates whether a mechanical abstraction of a task can induce an immediate improvement in bimanual multi-fingered performance. In a previous study on hemiparetic bimanual behavior (Itkonen et al., 2019), we observed that subjects mentally conceptualized two independently rotated wheels as a single unit. Here, we aim to further explore how conceptualization influences bimanual performance using a more quantifiable approach.

*Reaction time* (RT) is the duration between a stimulus and the initiation of an action. In a *simple reaction time* (SRT) task, the response is preplanned and triggered by an expected stimulus. In contrast, a *choice reaction time* (CRT) task requires additional cognitive processing, including stimulus discrimination, response selection, and motor programming, before movement initiation (Blinch et al., 2014).

In this study, we employed a response time test in which participants manipulated four symmetrically placed buttons in response to visual stimuli. In the first condition, participants pressed the buttons directly using their thumbs and index fingers. In the second condition, a plate was placed over the buttons, requiring participants to rock the plate to indirectly activate them. We hypothesized that this mechanical abstraction, introduced without verbal explanation, would prompt participants to form a new mental model of the task, thereby improving response times. Finally, we removed the plate to assess whether the conceptualization persisted after its removal.

## 2 Materials and methods

### 2.1 Subjects

Initially, 43 volunteers participated in the experiment. The sample size was based on similar studies measuring bimanual response times (Öttlet al., 2023; Blinch, Holmes, Cameron, & Chua, 2021). As the initial hypothesis was not confirmed in the full population, the data were analyzed in subgroups and supplemented with a control group (Table 1).

**Table 1:**
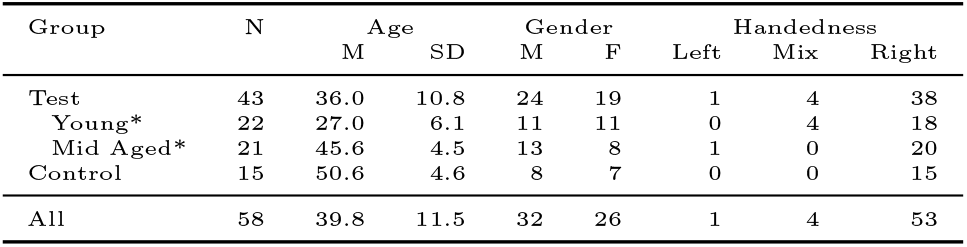
Summary of Subjects. The groups “Young” and “Middle-Aged”, marked with an asterisk (*), are subgroups within the Test group. “All” includes both Test and Control groups.

| Group | N | Age |  | Gender |  | Handedness |  |  |
| --- | --- | --- | --- | --- | --- | --- | --- | --- |
|  |  | M | SD | M | F | Left | Mix | Right |
| Test | 43 | 36.0 | 10.8 | 24 | 19 | 1 | 4 | 38 |
| Young* | 22 | 27.0 | 6.1 | 11 | 11 | 0 | 4 | 18 |
| Mid Aged* | 21 | 45.6 | 4.5 | 13 | 8 | 1 | 0 | 20 |
| Control | 15 | 50.6 | 4.6 | 8 | 7 | 0 | 0 | 15 |
| All | 58 | 39.8 | 11.5 | 32 | 26 | 1 | 4 | 53 |

To contrast younger and older participants, the test group was divided into two subgroups: “Young Adults” (20–39 years) and “Middle Aged” (40–59 years). This classification, commonly used in reaction time research (Dykiert, Der, Starr, & Deary, 2012), ensured an even split between age groups.

Following the initial experiment, an additional 15 participants were recruited as a control group to match the “Middle Aged” participants. A sample size of *N* = 15 was determined using G*Power 3.1.9.6 (Faul, Erdfelder, Lang, & Buchner, 2007) for a one-sample t-test assessing deviation from a known constant. Parameters included: Effect Size *d* = 0.7 (based on bimanual CRT changes in “Middle Aged”), *One T ail, α* = 0.05, and *Power* (1 *− β*) = 0.95.

Handedness was assessed using the short-form Edinburgh Handedness Inventory (Veale, 2014): Left = 1, Mixed = 4, Right = 38. Participants rated their handedness based on writing, throwing, toothbrush use, and spoon use, with responses scored as 100 for “Always” and 50 for “Usually.” An average score below 60 indicated “mixed handedness.”

The experiment was exempt from approval by the University of Eastern Finland Committee on Research Ethics, as it did not meet the criteria set by the Finnish Advisory Board on Research Integrity. The study complies with the Helsinki Declaration, and all participants provided written informed consent.

### 2.2 Device

A dedicated response device was designed for registering response times (see Figure 1b). The device comprised four momentary tactile push buttons (diameter: 6.5 mm, travel distance: 0.5 mm) arranged in a square formation. Each button required an activation force of approximately 200 grams and provided an audible click upon activation. The lateral and longitudinal distances between the centers of the buttons were both set to 50 mm. During the experiments, one finger (thumb or index) was assigned to each button. The buttons were wired to an Arduino-compatible microcontroller, which detected changes in button states and transmitted the data to the host computer via a serial connection. The host computer displayed stimuli, managed the experimental tasks, and logged the results.

**Figure 1:**
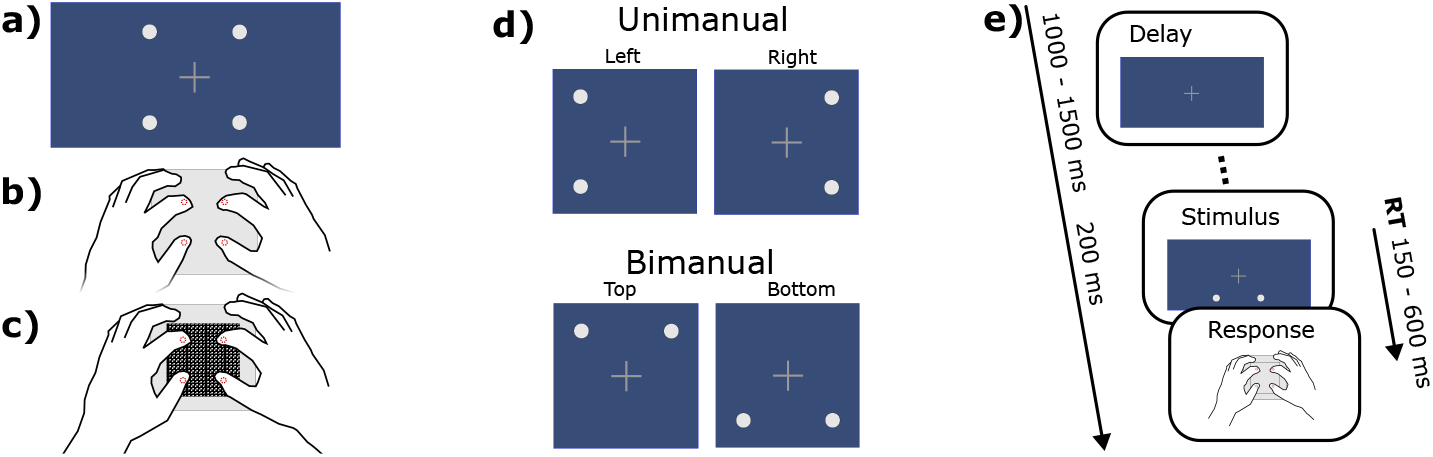
a) The stimuli are displayed on a computer screen with a static fixation cross and four disks that can be individually shown or hidden to create different stimulus variants. b) The response device has four buttons spatially aligned with the disks on the screen, with one finger assigned to each button, where fingers remain in position throughout the entire task duration. Dashed-line circles represent the buttons. c) The response device in the “plate” condition, where the check-patterned square depicts a plate placed between the fingers and the buttons. d) The four stimulus patterns for two-finger tasks include two unimanual tasks, activated by the index finger and thumb of the same hand, and two bimanual tasks, activated by the index fingers or thumbs of both hands. e) Each trial begins after a random delay, with the stimulus displayed for 200 ms. Response times (RT) are measured from stimulus onset.

### 2.3 Plate

To modulate changes in behavior, a 90*×*90 mm square plate was placed on top of the push buttons (see Figure 1c). The plate was constructed from two transparent 2 mm plastic sheets, one featuring 10 mm diameter holes aligned with the buttons. The plate weighed approximately 20 grams. Cavities on the underside stabilized it over the buttons, allowing it to rest freely. Pressing at the edge of the plate could cause it to fall off. The positions of the underlying buttons were marked with small glue bumps on the top surface, enabling subjects to position and maintain their fingers on the buttons without looking.

While using the plate, subjects mediated button presses by rocking it. However, plate activation required simultaneous pressure from two fingers. Subjects were instructed to press the buttons as before, now with the plate between their fingers and the buttons. This setup enabled four distinct two-finger interaction patterns (see Figure 1d), classified as unimanual or bimanual depending on whether activations involved one or both hands.

### 2.4 Stimuli and Response

Subjects were instructed to place their fingers on the buttons of the device and to fixate their gaze on a cross at the center of the computer display. Stimuli were presented as flashing discs arranged in a square constellation around the fixation cross (see Figure 1a). Before each task, the buttons were linked to on-screen feedback to help subjects establish an understanding of the stimulus-response mapping. A single trial is illustrated in Figure 1e).

Participants were instructed to respond as quickly as possible, disregarding any potential errors. To prevent prioritization of accuracy over speed, a response time limit of 600 ms was set. This limit was determined during pretesting as conservative enough to capture responses when participants were vigilant. Pretesting also revealed that providing feedback on late responses caused some participants to slow down, so no such feedback was given. If a response exceeded 600 ms or included incorrect buttons, the trial was added to the end of the task list for repetition.

### 2.5 Protocol

The core of the experiment was a four-choice reaction time (4-CRT) task, conducted under three conditions to assess the effect of the plate. Familiarization with the task and its conditions was achieved through simple reaction time (SRT) tasks. Figure 2 illustrates the experimental protocol, detailing the order of tasks alongside an example of registered data.

**Figure 2:**
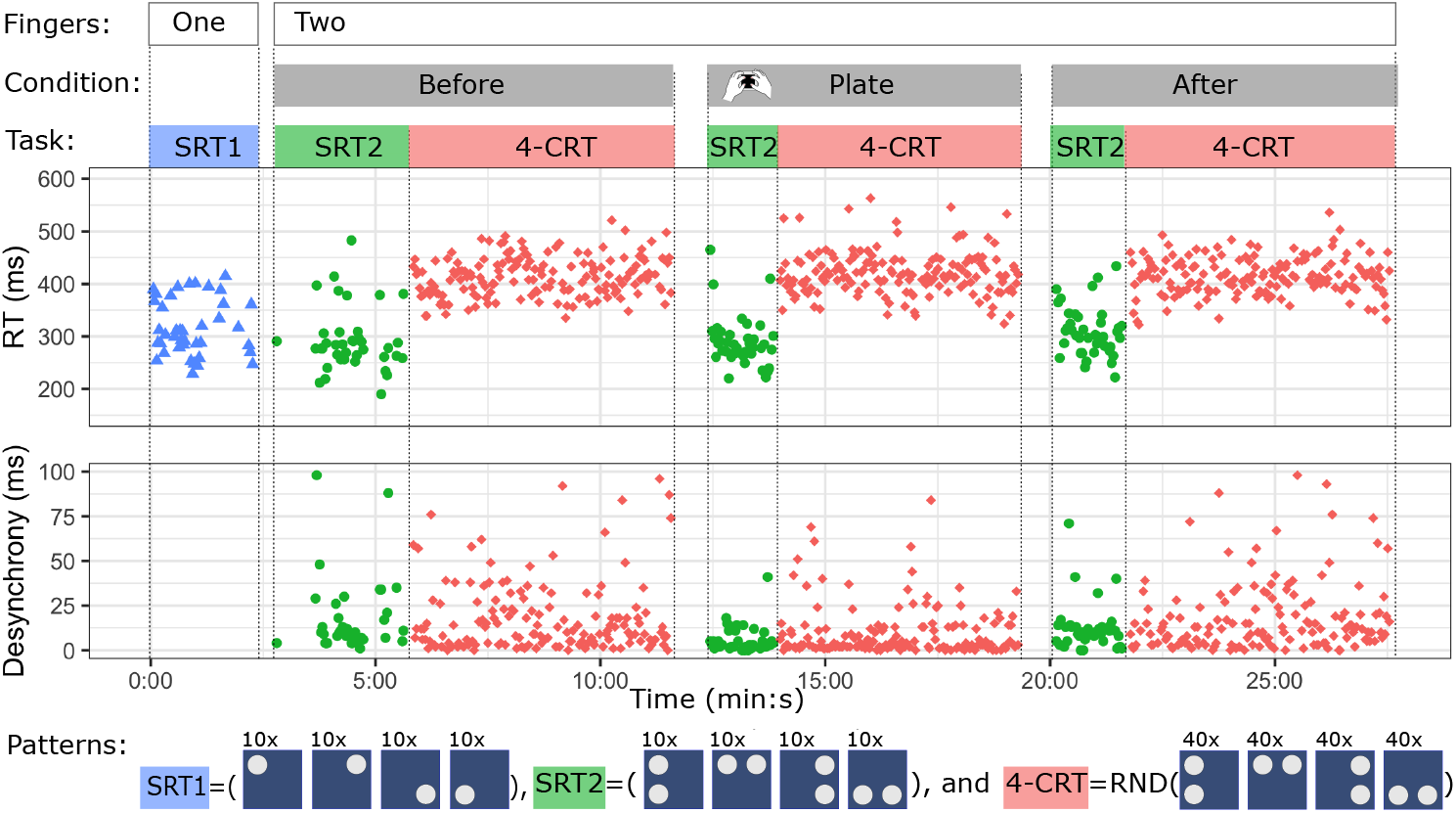
Full protocol sequence with example registered data. The session began with single-finger SRT1 tasks, consisting of 10 trials on each of the four device buttons (4 *×*10 trials, total 40 trials). The remaining tasks were two-finger tasks performed in three conditions: Before, Plate, and After. In the “Plate” condition, a plate mediated the response between the fingers and buttons (3*×* 4*×* 10 trials, total 120 trials). Each condition included SRT2 tasks with 10 repetitions of each two-finger pattern, and 4-CRT tasks with 40 repetitions of each two-finger pattern in randomized order (3*×* 4 *×*40 trials, total 480 trials). Registered data included reaction times (RT) and desynchrony, defined as the timing difference between the two-finger responses. The control group did not use the plate; instead, they completed the middle condition without it, in the same way as the other two conditions.

The experiment session began with a single-finger SRT task. This task consisted of 10 trials for each button in the following order: top-left, top-right, bottom-left, and bottom-right. Participants were asked whether they felt comfortable pressing the device with each finger and were monitored to ensure they could respond in a timely manner. They were also asked if they were ready before the initiation of each new task. Between tasks, finger placements were corrected as needed, and participants were encouraged to find a comfortable posture.

The remainder of the session included three conditions: *Before, Plate*, and *After*, each consisting only of two-finger trials. For the Test group, the plate was used only in the middle condition, while the Control group did not use the plate in any condition. Each condition began with an SRT task, followed by a CRT task. Each SRT task consisted of 10 trials in the following sequence: Left, Top, Right, Bottom. The CRT tasks included 40 randomized repetitions of each prior task. Randomization was conducted in batches of three cycles (12 trials) to ensure a balanced distribution of patterns.

After completing the session, participants assessed their experience with the device and plate by answering two questions: “Did you feel a difference between the trials using the additional plate and without the plate, on a scale from 1 (no difference) to 10 (totally different)” and “Which one did you prefer: with the plate or without the plate?”

### 2.6 Data registration

The data recording consisted of two types of response times: the reaction time and the desynchrony time. The *reaction time* (RT) was calculated as the difference between the time from the onset of the stimulus to the first activation of the target buttons. Valid responses required simultaneous activation of all target buttons. If any incorrect button was pressed, the trial was automatically rejected and repeated at the end of the sequence. Responses with reaction times shorter than 150 ms were rejected due to subject anticipation. The *desynchrony time* was calculated as the difference between the time of the first contact with the button and the time when both required buttons were pressed. Responses with desynchrony times longer than 100 ms were rejected. The average experiment session lasted 28 minutes and 40 seconds (SD = 3 minutes and 16 seconds).

The responses for each subject were aggregated as means over valid trials. Due to fundamental differences in biomechanics and neurophysiology, the two-finger responses in the SRT2 and 4-CRT tasks were grouped into bimanual and unimanual patterns, and the related reaction times were calculated as follows:

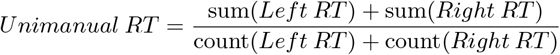

and

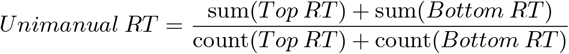

See Figure 1 d) for visual presentation of the patterns.

### 2.7 Data analysis

Data were initially analyzed for the Test group as a whole to compare responses between conditions and across bimanual and unimanual patterns. After confirming the null hypothesis, further exploration was conducted by splitting the population into two age-based groups for comparison, while age correlations were analyzed for the entire population. All analyses were performed using the same tools and methodology described below.

Data were analyzed using R version 4.4.1 (R Core Team, 2019). The “rstatix” package (Kassambara, 2023) version 0.7.2 was utilized to facilitate the implementation of statistical tests.

Statistical analyses comparing two distributions were conducted using a paired Student’s t-test. If normality was not met, as determined by the Shapiro-Wilk test (p *<* 0.05), a paired Wilcoxon signed-rank test was used instead.

For comparisons involving more than two distributions, repeated measures ANOVA was applied. Outliers exceeding three times the standard deviation were removed. When sphericity was violated (Mauchly’s test p *<* 0.05), the Greenhouse-Geisser correction was applied. Post hoc analyses were performed using pairwise t-tests with Bonferroni correction.

Comparisons of correlations were conducted using the “cocor” library (Diedenhofen & Musch, 2015). Significance was determined by 95% confidence intervals, using a comparison of two overlapping correlations based on dependent groups, applying Zou’s method (Zou, 2007).

### 2.8 Transparency and openness

We report how we determined our sample size, all data exclusions (if any), all manipulations, and all measures in the study, and we follow the Journal Article Reporting Standards (JARS) (Appelbaum et al., 2018). All data, analysis code, and research materials are available on request within the limits of General Data Protection Regulation (GPDR) and participants’ consents. This study’s design and its analysis were not pre-registered.

## 3 Results

In this study, we explored the impact of the introduction of the plate on behavior in a multi-fingered bimanual manipulation task. The analysis includes a characterization of tasks and patterns, followed by an examination of reaction times and desynchrony—the inter-digit temporal difference in multi-fingered activations—separately for bimanual and unimanual patterns. The hypothesis was partially supported, as the expected effect of reduced reaction times was observed only in bimanual patterns, not in unimanual ones. Age-related differences in the effect were highlighted through age-split group analyses and supported by correlations with age. To validate the primary result, the experiment was repeated with a control group that performed the task without using the plate.

### 3.1 Simple reactions

The results for the first task involving single-digit responses (SRT1) are summarized in Table 2. The sessions then progressed to two-finger tasks (SRT2) performed in four directions. However, due to the limited number of trials and the standardized task sequence across participants, rigorous comparisons between individual fingers or specific directions were not feasible.

**Table 2:**
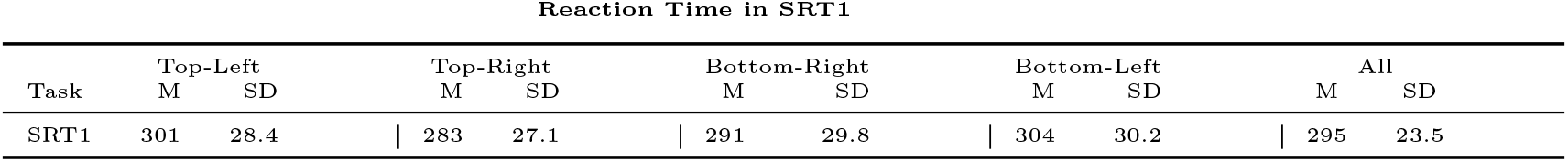
Reaction times (RT) in milliseconds (ms) for single-finger responses across all participants. The column “All” represents the mean of all responses for the four individual buttons. During the session, SRT1 was recorded only once with 10 trials on each button.

For patterns involving two fingers, participants were instructed to press buttons simultaneously. Consequently, the delay between activations (desynchrony) reflects the difficulty of executing the pattern. The mean SRT2 increased in the test group with the introduction of the plate by 15 ms (SD = 21.2; *t*(42) = *−*4.76, *p <* 0.001). This effect was more pronounced in unimanual responses (22 ms, SD = 17.2) compared to bimanual responses (8 ms, SD = 21.1). The plate decreased mean desynchrony by 7 ms (SD = 5.9; *t*(42) = *−*7.39, *p <* 0.001), which was again more pronounced in unimanual responses (9 ms, SD = 7.9) compared to bimanual responses (4 ms, SD = 7.7).

Additionally, two-finger simple reactions were 7 ms faster (SD = 19.3, t(57) = 2.91, p = 0.005) than single-finger responses.

### 3.2 Effects of hand and finger use on choice reactions

Following the simple reaction time tasks (SRT1 and SRT2), participants proceeded to the first 4-CRT task, which was repeated 40 times for each pattern. The initial measurements for these tasks, recorded before the plate interventions, were comparable across all participants, including the control group. These results are summarized alongside SRT2 in Table 3.

The ANOVA, calculated over all trials in the tasks, showed a significant difference in choice reaction times across patterns (F(2.00, 79.22) = 504.81, *p <* 0.001, *η*^2^ = 0.71). There was no significant difference between left and right sides (t(57) = 0.69, p = 0.496). However, responses to the top direction (index finger activations) were 11 ms faster (SD = 18.0) compared to the bottom direction (thumb activations). Desynchrony for thumb activations was 3 ms longer (SD = 7.9) compared to index fingers (W = 546, p = 0.017).

**Table 3:** Response times: reaction time (ms) and desynchrony (ms) during the first of three rounds of two-fingered patterns before the introduction of the plate. The column “All” represents the mean of responses across the four individual directions.

| Response Times in First SRT2 and First 4-CRT |  |  |  |  |  |  |  |  |  |  |  |  |  |  |  |  |  |  |  |  |
| --- | --- | --- | --- | --- | --- | --- | --- | --- | --- | --- | --- | --- | --- | --- | --- | --- | --- | --- | --- | --- |
|  | Left |  |  |  | Top |  |  |  | Right |  |  |  | Bottom |  |  |  | All |  |  |  |
|  | RT |  | Desync |  | RT |  | Desync |  | RT |  | Desync |  | RT |  | Desync |  | RT |  | Desync |  |
|  | M | SD | M | SD | M | SD | M | SD | M | SD | M | SD | M | SD | M | SD | M | SD | M | SD |
| SRT2 | 297 | 38.5 | 23 | 10.0 | 279 | 32.6 | 17 | 10.2 | 286 | 32.8 | 21 | 9.7 | 290 | 37.4 | 18 | 10.3 | 288 | 31.2 | 20 | 6.2 |
| 4-CRT | 395 | 42.8 | 29 | 9.2 | 405 | 50.4 | 16 | 5.9 | 393 | 40.4 | 26 | 8.7 | 416 | 49.0 | 18 | 7.5 | 402 | 44.1 | 22 | 5.9 |

In contrast to the findings from the simple reaction tasks, unimanual responses in the first choice reaction tasks were 16 ms faster than bimanual responses (SD = 16.5, t(57) = *−*7.61, p *<* 0.001). This trend was reversed in the simple two-finger responses, where unimanual reactions were 7 ms slower than bimanual responses (SD = 18.8, t(57) = *−*2.86, p = 0.006).

### 3.2 Desynchrony in choice reactions

The plate physically connects the fingers that activate the buttons. As a result, during plate use, desynchrony times in the 4-CRT task decreased for both bimanual and unimanual activations (see Figure 3). This change did not correlate with age (r(41) = 0.17, p = 0.275). The desynchrony measurements in the 4-CRT task are summarized in Table 4.

**Table 4:**
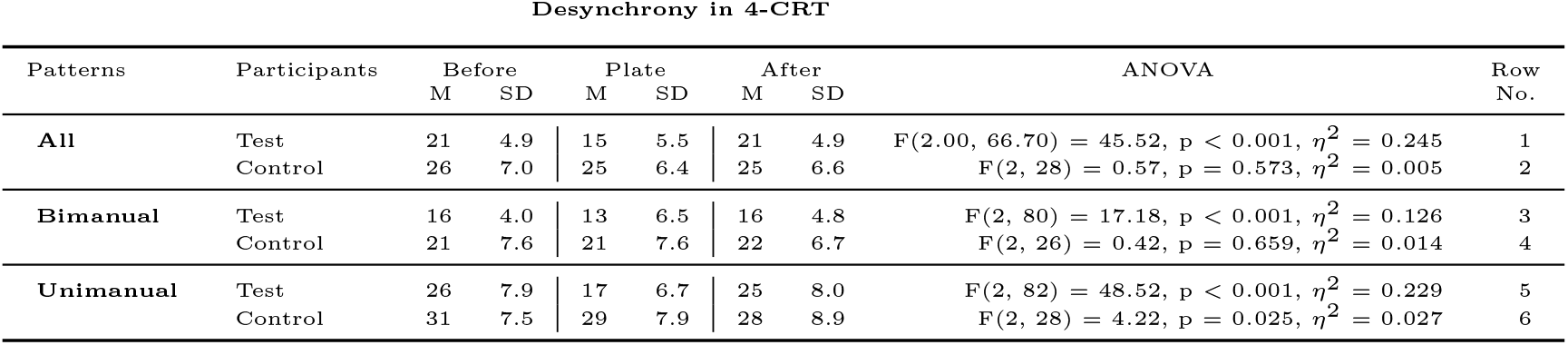
Desynchrony times in milliseconds (ms) for the Before, Plate, and After conditions. A within-subjects repeated measures ANOVA was conducted to compare desynchrony times across the conditions.

**Figure 3:**
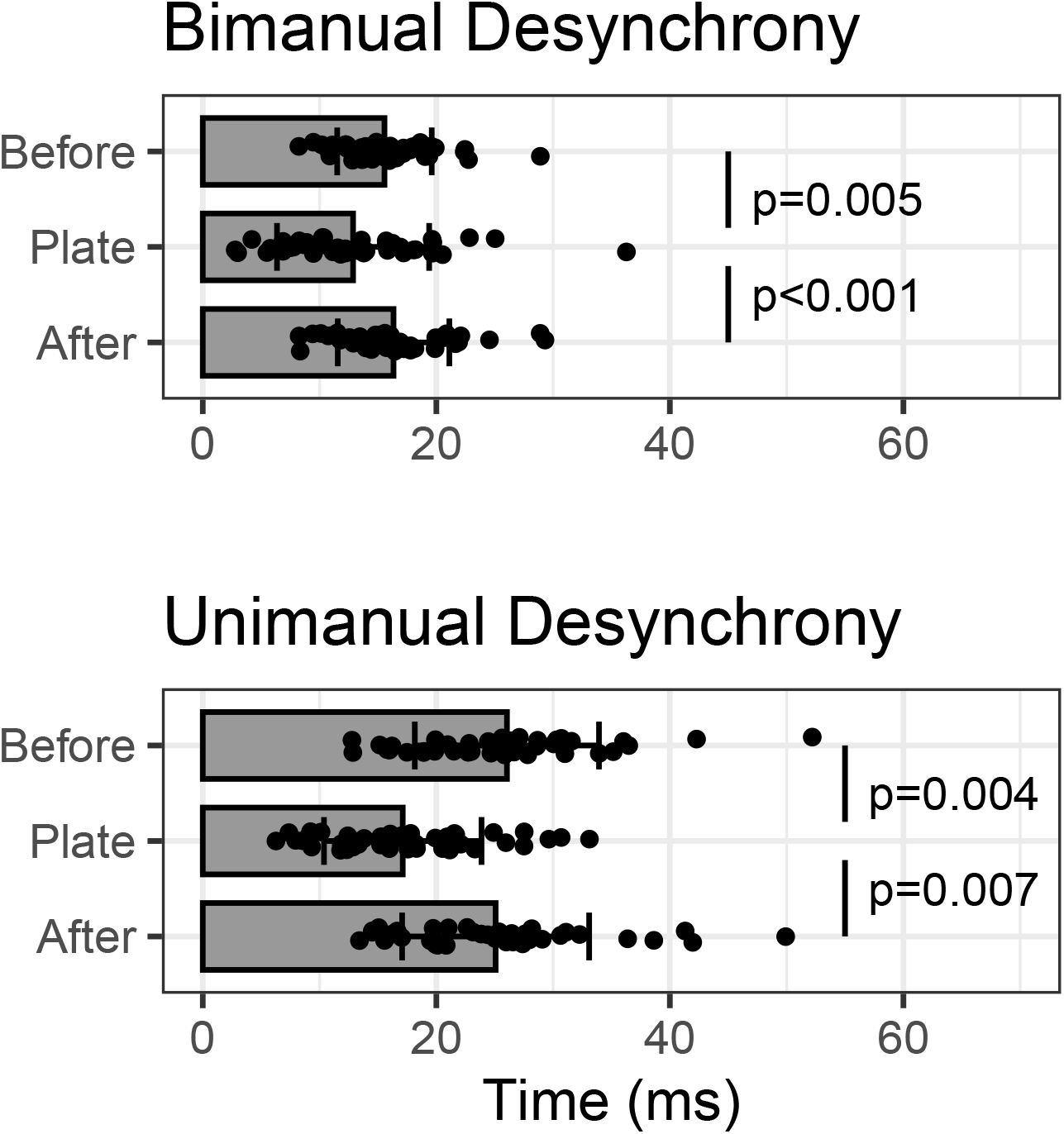
Desynchrony times in 4-CRT responses for the Test group participants. The time difference between finger presses decreased with the use of the plate. The whiskers indicate *±*SD. ANOVAs and values are in Table 4.

The average reduction in desynchrony from the Before condition to the Plate condition was 3 ms (SD = 5.3) for bimanual patterns and 9 ms (SD = 7.6) for unimanual patterns. Desynchrony times between the Before and After conditions did not differ significantly.

The control group performed the protocol by repeating the 4-CRT task three times, all without the plate. According to the ANOVA, there was no significant change in desynchronization for the bimanual patterns (Table 4, row 4). For unimanual patterns, the ANOVA indicated significant variability across the three rounds, but post-hoc analysis did not confirm any significant differences. The observed 3 ms (SD = 4.6) decrease in desynchrony from Before to After condition was not statistically significant.

### 3.4 Reaction time in choice reactions

Contrary to our hypothesis, reaction times (RT) in the 4-CRT task did not show a significant change from the Before to After condition (see Table 5, row 1). However, a significant change was observed in bimanual patterns (row 5), whereas unimanual patterns did not exhibit such a change (row 9).

**Table 5:** Reaction time (RT) in milliseconds (ms) for the Before, Plate, and After conditions, with changes between the Before and After conditions grouped by patterns and participants. The test statistic compares reaction times between the Before and After conditions, using a paired t-test when the data follow a normal distribution; otherwise, the Wilcoxon signed-rank test is used. The participants ‘Young” and “Middle-Aged,” marked with an asterisk (*), are subgroups within the Test participant group.

| Reaction Time in 4-CRT |  |  |  |  |  |  |  |  |  |  |  |
| --- | --- | --- | --- | --- | --- | --- | --- | --- | --- | --- | --- |
| Patterns | Participants | Before |  | Plate |  | After |  | Change |  | Before vs. After | Row No. |
|  |  | M | SD | M | SD | M | SD | M | SD | Test Statistic |  |
| All | Test | 396 | 45.3 | 398 | 40.8 | 390 | 40.2 | -5 | 19.4 | $t(42) = 1.82, p = 0.076$ | 1 |
| | Young* | 379 | 44.0 | 389 | 44.4 | 379 | 42.8 | 0 | 16.4 | $t(21) = -0.02, p = 0.983$ | 2 |
| | Mid Aged* | 413 | 40.3 | 408 | 35.2 | 402 | 34.2 | -11 | 21.0 | $t(20) = 2.42, p = 0.025$ | 3 |
| | Control | 422 | 35.1 | 423 | 40.3 | 422 | 38.3 | 1 | 19.8 | $t(14) = -0.12, p = 0.910$ | 4 |
| Bimanual | Test | 403 | 50.4 | 400 | 45.2 | 394 | 44.4 | -8 | 21.3 | $t(42) = 2.62, p = 0.012$ | 5 |
| | Young* | 382 | 48.2 | 390 | 49.3 | 381 | 45.4 | -2 | 19.5 | $t(21) = 0.42, p = 0.676$ | 6 |
| | Mid Aged* | 424 | 44.5 | 411 | 38.8 | 408 | 39.6 | -16 | 21.2 | $t(20) = 3.37, p = 0.003$ | 7 |
| | Control | 433 | 36.7 | 433 | 39.3 | 434 | 38.7 | 1 | 21.4 | $t(14) = -0.11, p = 0.915$ | 8 |
| Unimanual | Test | 388 | 41.1 | 395 | 37.5 | 386 | 37.3 | -2 | 19.3 | $t(42) = 0.70, p = 0.488$ | 9 |
| | Young* | 375 | 40.6 | 387 | 40.2 | 377 | 41.2 | 2 | 16.0 | $t(21) = -0.58, p = 0.566$ | 10 |
| | Mid Aged* | 403 | 37.5 | 404 | 33.2 | 396 | 30.6 | -6 | 21.8 | $W = 167, p = 0.076$ | 11 |
| | Control | 410 | 35.3 | 413 | 43.3 | 411 | 39.2 | 1 | 20.2 | $W = 62, p = 0.934$ | 12 |

To further investigate these differences, participants were divided into two subgroups based on age. Figure 4 illustrates RT in bimanual and unimanual responses for these groups. Among the subgroups, bimanual patterns in middle-aged participants were the only group to show a significant change between the Before and After conditions (Table 5, row 7). This finding suggests that the plate intervention specifically improved RT in this group. In the case of young participants, the plate did not improve response times, and their mean response time with the plate was slower.

The split at 40 years of age is arbitrary as the age related changes in nervous system take place gradually. The reaction times were investigated in relation to age by means of correlation analysis. Figure 5 illustrates the correlation of CRT in bimanual and unimanuial patterns with age. The bimanual correlation dropped from the Before condition r(41) = 0.53, p *<* 0.001 to r(41) = 0.37, p = 0.014 in the Plate condition and remained at r(41) = 0.44, p = 0.003 in the After condition. Comparison of correlations showed statistical difference between the Before and Plate conditions (Zou’s 95% confidence interval for r.jk - r.jh: 0.0079 0.3235). The corresponding values for correlation of unimanual CRT with age was Before (r(41) = 0.42, p = 0.005), Plate (r(41) = 0.38, p = 0.013), and After (r(41) = 0.40, p = 0.008).

**Figure 4:**
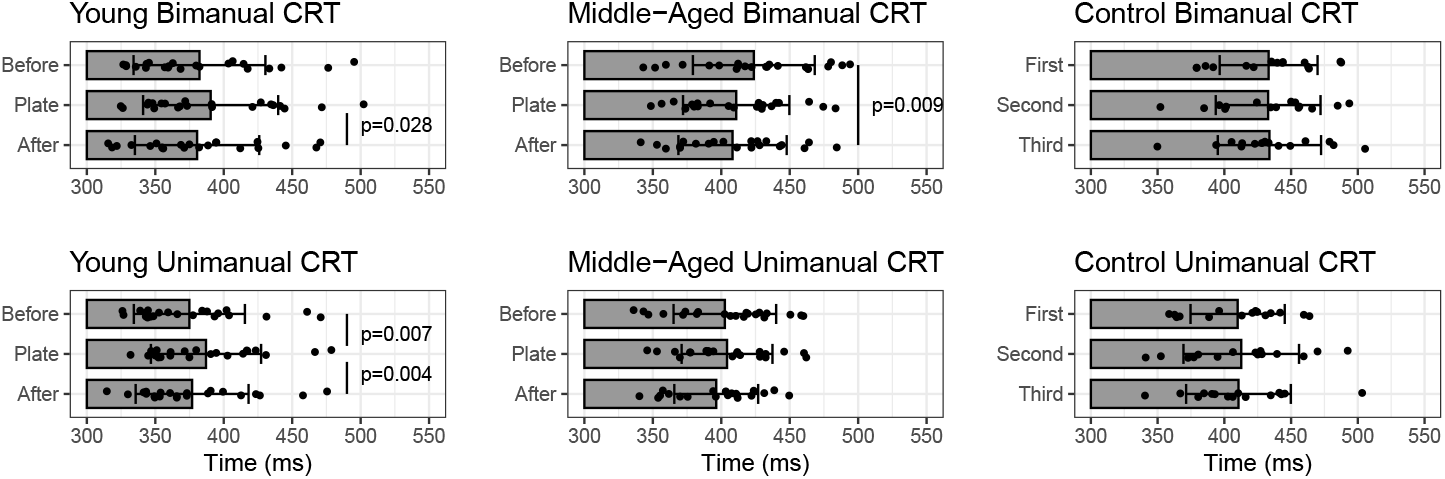
Bimanual and unimanual CRT tasks in the Young (20–39 years) and Middle-aged (40–59 years) groups, as well as the control group. Significant changes in bimanual CRT were observed in the Middle-aged group following the plate intervention. Significant differences are marked with an asterisk (*), while other differences are not statistically significant. Whiskers represent *±*SD. ANOVAs are Young Bimanual: F(2, 42) = 3.48, p = 0.040, *η*^2^ = 0.009; Young Unimanual: F(2, 42) = 7.99, p = 0.001, *η*^2^ = 0.018; Middle-Aged bimanual:F(1.38, 27.60) = 6.36, p = 0.011, *η*^2^ = 0.028; Middle-Aged bimanual: F(2, 40) = 1.69, p = 0.198, *η*^2^ = 0.011, Control Bimanual: F(2, 28) = 0.977, p = 0.98, *η*^2^ = 0.0001; and Control Unimanual: F(2, 28) = 0.157, p = 0.86, *η*^2^ = 0.0009.

**Figure 5:**
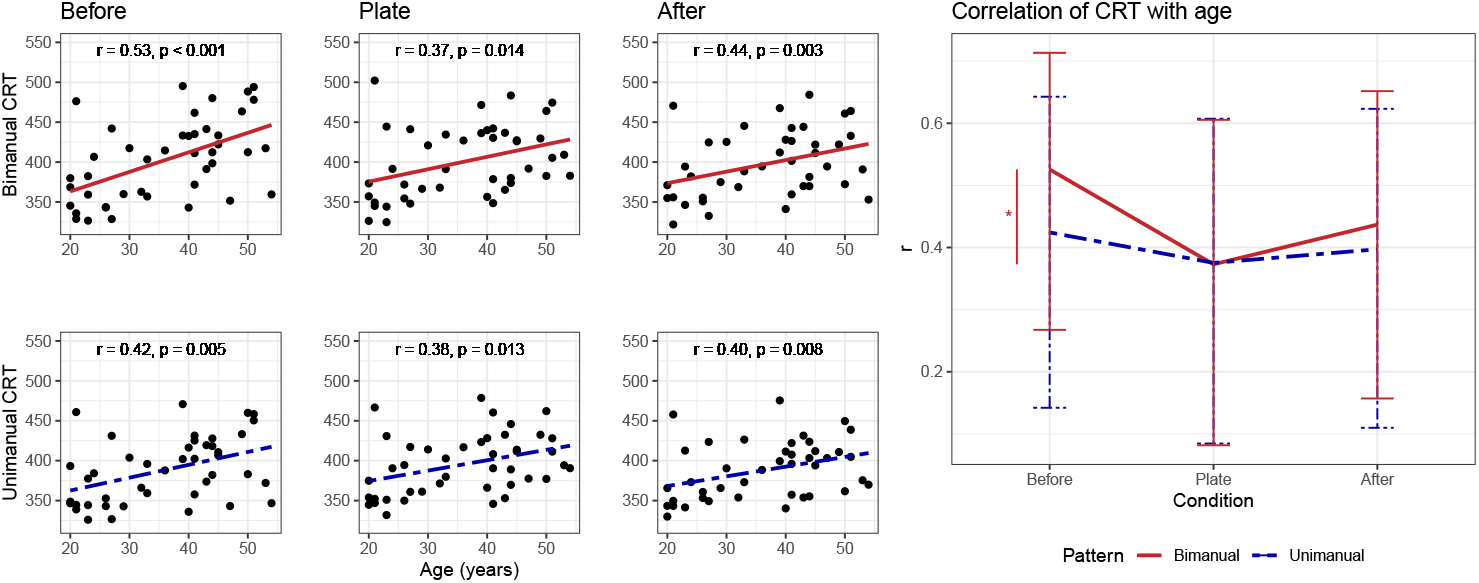
Correlation of 4-CRT with age under each condition: The correlation of bimanual CRT with age is significantly reduced from Before condition to Plate condition. Conversely, the change in unimanual correlation with age is not significant. “r” represents Pearson’s coefficient of correlation. The whiskers mark 95% confidence interval.

### 3.5 Control study

As previous results suggested that the plate intervention led to age-related improvement in bimanual reaction times, this improvement could also be attributed to learning effects. To address this, an additional control group, matching the middle-aged portion of the Test group, was recruited. This group followed the same protocol as the Test group but without using the plate, instead repeating the same SRT2 and 4-CRT routine three times. Desynchrony in bimanual patterns did not change (Table 4, row 4). While ANOVA for unimanual patterns showed significance (Table 5, row 6), corrected post-hoc values did not reveal significant interactions. Reaction times did not indicate a change from the Before to After condition (Table 5, rows 8 and 12). The reaction times for the control group are shown in Figure 4.

In the controlled study, the middle-aged portion of the Test group was compared against the Control group. A repeated-measures ANOVA was conducted to detect changes in reaction time from the Before to After condition in bimanual patterns for the Test group but not in unimanual patterns, with no change expected in the Control group. The within-subject factors were Condition (Before, After) and Pattern (Bimanual, Unimanual), while Group (Test, Control) served as the between-subject factor. The hypotheses tested included the effect of condition on reaction time, the effect of pattern on reaction time, and the interactions between condition, pattern, and group.

Repeated-Measures ANOVA results revealed a significant main effect of Pattern on reaction time, F(1,34) = 61.028, p *<* 0.001, *η*^2^ =0.066, indicating faster reaction times in unimanual patterns compared to bimanual patterns. A significant Condition–Pattern interaction, F(1,34) = 6.14, p =0.018, *η*^2^ =0.001, suggesting that the effect of condition on reaction time differed between bimanual and unimanual patterns. A significant Condition–Pattern–Group interaction, F(1,34) = 6.09, p =0.019, *η*^2^ = 0.001, supporting the hypothesis that the Test group exhibited a change in reaction times for bimanual patterns but not for unimanual patterns, whereas no significant changes were observed in the Control group. No other main effects or interactions reached statistical significance.

Post hoc analyses revealed a significant difference in reaction times from the Before to After condition for the Test group in bimanual patterns (t(20) = 3.37, p = 0.005). No significant differences were observed for the Test group in unimanual patterns (t(20) = 1.32, p = 0.2), or for the Control group in either bimanual (t(14) = *−*0.109, p = 0.915) or unimanual patterns (t(14) = *−*0.119, p = 0.907).

### 3.6 Response accuracy

The valid responses were those that were correct and accepted within the time range between the stimulus presentation and the maximum allowed response time. To evaluate participant performance and task compliance, we calculated the hit rate, defined as the ratio of accepted responses to presented stimuli. Table 6 summarizes the ratios of valid responses for all participants.

**Table 6:**
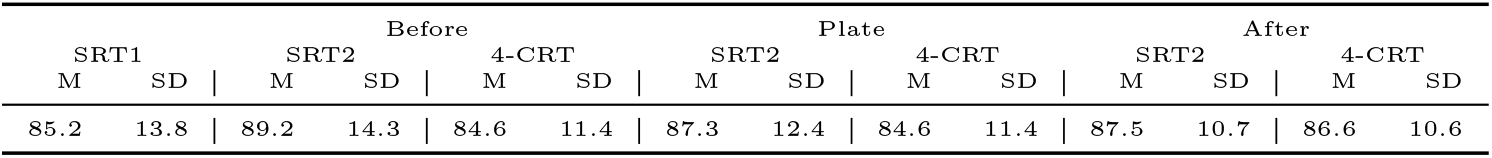
The percentages of valid responses in all tasks and conditions across all participants.

### 3.7 Perceived Difference for Plate Intervention

After completing the experiment, participants evaluated whether they perceived a difference between tasks performed with and without the plate, using a scale from 1 (no difference) to 10 (completely different). The mean difference rating was 6.5 (SD = 1.6), with no significant correlation with age (r(41) = 0.1, p = 0.5).

A follow-up question assessed participants’ preference for either interaction mode. Overall, 24 participants (56%) preferred using the plate. Among young participants, 45% preferred manipulation with the plate, compared to 67% of middle-aged participants who expressed the same preference.

## 4 Discussion

When designing the experiment, we hypothesized that augmenting the four-button interface with a plate would implicitly shift the mental representation of the task from pressing two buttons to rocking a plate in one direction. However, our analysis revealed that this effect was not observed when considering the sample as a whole, contrary to our initial hypothesis. Further analysis showed an effect in bimanual tasks, these were the tasks where both hands acted symmetrically. Notably, this effect was absent in younger subjects, whose responses remained unaffected by the plate intervention. These findings suggest that the impact of the plate on task performance is age-dependent.

### 4.1 Facilitation of bimanual responses

We observed that the plate facilitated bimanual behaviors in middle-aged subjects, whereas unimanual behaviors showed no significant improvement. To understand this effect, we first examine the underlying mechanics of bimanual control.

Bimanual control is primarily governed by the sensorimotor cortices, which control contralateral arms (Purves, Augustine, Fitzpatrick, Hall, & LaMantia, 2018). The arms also reflect signals to the ipsilateral sensorimotor cortices: stimulation of these cortical areas induces limb movements, while sensory stimulation of an arm elicits cortical responses. The cortices are interconnected via the corpus callosum, which contains both inhibitory and excitatory pathways (Swinnen, 2002).

Both unimanual and bimanual actions activate cortical regions in both hemispheres, but these interactions are complex enough that bimanual cortical activations are not necessarily stronger than unimanual ones (Koeneke, Lutz, Wüstenberg, & Jäncke, 2004; Derosière et al., 2014). Interhemispheric inhibition in the primary motor cortex plays a key role in bimanual movement control (Morishita, Timmermann, Schulz, & Hummel, 2022; Hinder, Schmidt, Garry, & Summers, 2010). Increasing evidence of the participation of both cortical hemispheres in unilateral motion has led some researchers to speculate that unimanual movements may be a specific case of bimanual movements, where the opposing arm is actively inhibited (Chettouf, Rueda-Delgado, de Vries, Ritter, & Daffertshofer, 2020; Beaulé, Tremblay, & Théoret, 2012).

The bimanual motion patterns applied in the experiment were particularly easy to execute. Bimanual movements where contralateral homologous muscles contract simultaneously are more stable compared to other kinds of bimanual patterns (Haken et al., 1985). The overflow of motor commands to involuntary synchronous contractions of the opposite limb is known as *mirror movements*(Cincotta & Ziemann, 2008). These movements are observed in childhood, but they diminish as motor control develops. Weaker mirrored muscle activations, which do not necessarily result in motion, can also be observed in healthy subjects using surface electromyography(Cincotta & Ziemann, 2008). Task demand, fatigue, and age are factors that influence these movements (Bodwell, Mahurin, Waddle, Price, & Cramer, 2003). However, older individuals have been shown to inhibit mirror behaviors when they are aware of them (Addamo, Farrow, Bradshaw, Moss, & Karistianis, 2010).

Bimanual reactions were faster than unimanual ones in simple response tasks, supporting the role of interhemispheric inhibition, where unimanual actions incur an additional cost due to the need to suppress activation in the opposite limb. However, in contrast, this mechanism did not manifest in choice reaction tasks, indicating the involvement of higher-order cognitive processes. In our study, CRT tasks included a mix of bimanual and unimanual trials, yet bimanual responses did not interfere with unimanual ones. Given the ease of bimanual motion patterns, bimanual responses should have been faster, but our results showed the opposite. This suggests that task complexity and cognitive demands outweighed the natural efficiency of bimanual coordination, leading to slower reaction times.

### 4.2 Role of aging

The facilitation of bimanual behaviors was observed only in older participants. We demonstrated this by dividing participants into two groups, “Young” and “Middle aged,” and by analyzing the correlation between response time and age. Aging affects the nervous system, leading to slower response times, an effect more pronounced in CRT than in SRT (Woods, Wyma, Yund, Herron, & Reed, 2015). In our study, only CRT significantly correlated with age, whereas SRT remained unaffected. Since visual stimuli and motor execution were identical in both tasks, this suggests that age-related slowing primarily impacts cognitive and sensorimotor integration processes rather than basic perceptual or motor execution speed. This distinction is critical, as it supports the idea that the observed differences arise from changes in afferent somatosensory signals and cognitive preparation rather than from delays in visual processing or motor output.

Age-related deterioration of touch receptors and neural transmission is a gradual process in the peripheral nervous system (Kalisch, Kattenstroth, Kowalewski, Tegenthoff, & Dinse, 2012; McIntyre, Nagi, McGlone, & Olausson, 2021), leading to reduced tactile acuity (Reuter, Voelcker-Rehage, Vieluf, & Godde, 2012). Noisy and sometimes incongruent sensory cues are integrated within the central nervous system to form a coherent percept. Over time, the nervous system becomes more permissive in processing these signals (Mozolic, Hugenschmidt, Peiffer, & Laurienti, 2012), causing older individuals to perceive the same stimuli differently than younger individuals (Landelle, El Ahmadi, & Kavounoudias, 2018; DeLoss, Pierce, & Andersen, 2013). These changes in sensory integration may contribute to the increased variability and adaptation observed in older participants’ responses.

A manifestation of age-related changes in sensory processing can be observed in how haptic stimuli impact coordination. Serrien et al. (Serrien, Teasdale, Bard, & Fleury, 1996) found that haptic stimuli disrupted bimanual coordination in younger participants but not in older ones. In our study, the plate intervention caused a slowdown in response times for young participants, which may reflect their awareness of the intervention, as indicated by their post-experiment feedback. These findings suggest that younger participants’ response to plate-mediated haptic stimulation was slowed, possibly due to age-related differences in sensory processing.

### 4.3 Temporal aspects of the effect

The change in response time appeared after the introduction of the plate and also remained after its removal. As demonstrated by our control study, there was insufficient time for motor or sensory learning to occur. Instead, the change must have taken place in response generation control. Participants were not given any description of altered mechanical properties; they were simply instructed to press the buttons as before, now with the plate between their fingers and the buttons. Visual stimuli remained unchanged, ensuring that the only difference was the introduction of the plate, which altered sensory signals from the hands to the CNS.

We speculate that the plate intervention produced highly similar tactile and kinesthetic afferent signals from both hands. For older participants, this increased similarity in bimanual sensory input may have led to a different interpretation of the task, as haptic sensations were integrated within the CNS (Sohn, Niu, & Sanger, 2016). However, younger participants, with higher sensory acuity and better bimanual control, were not affected by this intervention.

### 4.4 Task conceptualization

The initial findings emerged while preparing for a study aimed at improving multi-fingered movements by using mental abstractions to manage high degrees of freedom in manipulation tasks. A four-button manipulandum was designed as a minimalistic device to provide a foundational understanding of multi-fingered interactions. Initial pilot testing involved participants from what would later be categorized as the middle-aged group. Improvement in reaction times following the removal of the plate used during the intervention occurred spontaneously, without explicitly introducing the conceptualization of the plate.

In this study, the manipulation protocol was repeated to achieve reproducible results with a reasonable number of participants. Replacing the physical intervention with a verbal or visual explanation was deferred for future research.

While we did not explicitly facilitate plate conceptualization, doing so might have positively influenced performance, as conceptualization has been successfully used to reduce bimanual interference. This has been achieved through instruction (Blinch et al., 2021) to guide response preparation or by modulating stimuli (Mechsner, Kerzel, Knoblich, & Prinz, 2001; Spencer, Ivry, Hazeltine, & Semjen, 2004; Denyer & Boyd, 2025).

We chose to keep stimuli constant across conditions to ensure that any observed differences could be attributed specifically to the effects of conceptualization rather than to the processing of variable visual stimuli. However, the unnecessary cognitive cost of translating visual stimuli from a computer display to the response device could be reduced with a stimulus display integrated into the response device (Blinch, Gooch, Clark, Murrin, & Bayouth, 2025). Additionally, the tasks in this study did not include asynchronous elements, which might have been more suitable for modulation through conceptualization, as seen in previous studies that primarily utilized asynchronous patterns.

### 4.5 Implications and future directions

The results showing facilitation only in bimanual responses and in middle-aged subjects, along with bimanual control strongly presented in the neurophysiology of brain structure and function, support the idea that conceptualization did not occur along the unimanual axis (i.e., between the fingers of the same hand). The absence of observed changes in simple responses suggests that more complex neural processes, such as motion selection, are involved. Future studies in task conceptualization should focus on bimanual interactions with increased complexity, with a particular consideration of age-related factors.

Modifications to the response device could allow for more detailed investigation of split-plate designs, enabling the haptic stimuli to be distributed either only intermanually or only intramanually. Intermanual connectivity could be achieved by placing one plate between the index fingers and another between the thumbs, while intramanual connectivity could be implemented by having one plate for the left hand and another for the right hand. Additionally, diagonal connectivity between the thumbs of opposing hands could provide asynchronous movement patterns, which were not explored in the current study due to the limitations of the response device.

A more detailed manipulation of the mechanical connection delivering haptic feedback between the fingers could improve our understanding of how haptic signals influence behavior. This influence may manifest as facilitation in older individuals or as a disturbance in younger ones. An extended study focusing on bimanual behaviors could further clarify these effects. In addition to transferring stimuli on mechanical media (the plate), tactile displays or proprioceptive stimuli could provide controlled ways to modulate sensory input in relation to bimanual actions.

The analysis of response times provides valuable insights into the cognitive demands of the control processes. However, more detailed neurophysiological measurements, such as corticomuscular coherence (Liu, Sheng, & Liu, 2019), would offer a deeper understanding of the dynamics in functional connectivity driving these processes.

## 5 Conclusions

We presumed that a mechanical augmentation would influence the behaviors of subjects. The influence was observed in the mode of synchronous bilateral movement patterns of middle-aged subjects. The effect of the augmentation was observable even after the mechanism was removed. The persistence of the changed behavior is yet to be determined.

The specific bimanual synchronous movement pattern is associated with undeveloped and also with degenerated motor performance. However, because we did not observe adverse influences in the unilateral pattern, we consider the observed effect as an improvement in performance. The variety of applied bimanual behaviors is far more complex, and it would be valuable to explore if other modes of bimanual behaviors could be augmented with a similar mechanism.

## 6 Author contributions

MI Conceived and designed research, MI & NL Performed experiments, MI Analyzed data, MI & RB Interpreted results of experiments, MI Prepared figures, MI Drafted manuscript, all authors edited and revised manuscript, and all authors approved final version of manuscript.

## Supporting information

Additional supporting information may be found in the online version of the article at the publisher’s website.

